# Epithelial stem cell-derived chemokines track clinical remission in inflammatory bowel disease in a disease-specific manner

**DOI:** 10.64898/2026.05.23.727433

**Authors:** Sanmi Alake, Anil Kadam, Traci Jester, Craig L. Maynard, Babajide A. Ojo

**Affiliations:** Department of Pediatrics, Division of Gastroenterology, Hepatology and Nutrition, University of Alabama at Birmingham, Birmingham, AL, USA; Division of Pulmonary, Allergy and Critical Care Medicine, University of Alabama at Birmingham, Birmingham, AL, USA; Gregory Fleming James Cystic Fibrosis Research Center, Birmingham, AL, USA; Department of Pathology, Division of Molecular and Cellular Pathology, University of Alabama at Birmingham, Birmingham, AL, USA; Department of Medicine, Division of Gastroenterology and Hepatology, University of Alabama at Birmingham, Birmingham, AL, USA

## Abstract

**Background and Aims:** Stem cell-derived organoids are promising platforms for therapeutic screening in inflammatory bowel disease (IBD), but identifying functional organoid readouts with translational utility is challenging. Colon epithelial organoids from patients with ulcerative colitis (UC) overexpress chemokines CXCL1, CXCL11, CCL2, and CCL28, yet whether these inflammatory signatures correlate with disease activity and treatment response is unknown. This short report investigates whether organoid-retained chemokines correlate with disease activity and therapeutic outcomes.

**Methods:** We interrogated three bulk and two single-cell transcriptomic datasets from IBD clinical trials encompassing anti-TNFα and anti-integrin therapies to determine whether epithelial chemokines retained in UC organoids track clinical response and distinguish treatment responders from non-responders to biologic therapy across multiple IBD patient cohorts.

**Results:** In bulk transcriptomic data, CXCL1, CXCL11, and CCL2 were elevated in active UC and normalized only in patients achieving clinical remission, independent of therapy class, with persistent chemokine overexpression in non-responders. Single-cell analysis demonstrated widespread chemokine overexpression in UC epithelial clusters, with consistent normalization of CXCL1, CXCL11, and CCL28 in LGR5-positive stem compartment of patients who achieved clinical remission, but not in non-responders. In Crohn’s disease, the resolution of these epithelial chemokines was not associated with clinical response.

**Conclusions:** Epithelial chemokines, particularly CXCL1, CXCL11, and CCL28, track clinical remission in UC and represent candidate biomarkers and functional endpoints for epithelial-directed therapeutic strategies using stem cell-derived UC organoid models.

## 1. Introduction

More than half of patients with inflammatory bowel disease (IBD) fail to achieve clinical remission on conventional immunotherapies ^1^. Since epithelial barrier integrity and mucosal healing are essential for sustained remission, identifying epithelial-directed therapeutic targets represents an unmet need. Patient-derived organoids (PDOs) have emerged as physiologically relevant platforms for preclinical drug screening in IBD ^2^; however, identifying functional endpoints in IBD organoids that are associated with therapeutic response and clinical remission is unclear.

A defining feature of IBD is the chemokine-directed leukocyte recruitment to the intestinal mucosa to propagate inflammation ^3^. In particular, cryptitis frequently precedes crypt abscess formation and epithelial barrier damage ^4, 5^. Notably, epithelial organoids from patients with ulcerative colitis (UC) showed retained overexpression of chemokines, including CXCL1, CXCL11, CCL2, and CCL28, which coordinate the recruitment of neutrophils, T cells, monocytes, and plasma cells ^6^. Thus, these epithelial chemokines may constitute a durable epithelial inflammatory program that may be therapeutically targeted in UC organoids. However, the clinical relevance of these epithelial chemokine signatures has not been fully established.

In this study, we analyze bulk and single-cell transcriptomic datasets from multiple clinical trials to determine whether epithelial chemokine expression reflects disease activity and therapeutic response in UC and CD.

## 2. Case Report

### 2.1 Methods

Publicly available transcriptomic datasets from clinical trials of biologic therapies in inflammatory bowel disease were analyzed, including anti-TNF (golimumab-GSE92415, infliximab-GSE73661) and anti-integrin (vedolizumab-GSE73661) cohorts with paired gene expression and clinical outcome data ^7-9^.

For bulk microarray datasets, data were extracted from GEO and analyzed in R (v.4.4.0). The Robust Multiarray Average method was used to obtain log2 expression values for each gene. log2-normalized probe intensities were analyzed using ANOVA with FDR adjustment. TPM data were analyzed with Mann-Whitney test for two groups, while Kruskal-Wallis test with Dunn’s post hoc was used for more than 2 groups. Statistical analyses were done with GraphPad Prism (v7.04). The expression of our chemokines of interest (CXCL1, CXCL11, CCL2, and CCL28) and Mayo scores in responders and non-responders were extracted for further analyses. Principal coordinate variable plot was developed using the factoextra package in R.

To view the expression pattern of chemokines of interest in the epithelium of IBD tissues, we analyzed the public single-cell RNA-seq data obtained from 18 UC patients and 12 healthy controls ^10^, single cell portal accession number: SCP259). Matrices and metadata for 123006 colon epithelial cells were downloaded from the UCSC Cell Browser (https://human-colon.cells.ucsc.edu), normalized with Seurat (v5.3.1), and visualized using the ggplot2 (v4.0.0) package in R.

To determine the cell type-specific resolution of epithelial chemokines following biologic therapy, publicly available single-cell transcriptomics data from the TAURUS-IBD study ^9^ was downloaded from Zenodo (https://doi.org/10.5281/zenodo.13768607), analyzed using Scanpy (v1.9.8), and visualized with Matplotlib (v3.9.2) in Python (v3.9.23). Mean log-normalized expression was compared to healthy controls using the Mann-Whitney U test. Multiple testing correction was performed using the Benjamini-Hochberg false discovery rate (FDR) method. Differences were considered statistically significant at |Log_2_ FC| ≥ 0.5 with FDR-adjusted p < 0.05.

### 2.2 Results

To investigate whether chemokines elevated in UC organoids exhibit a similar expression pattern and resolve in clinical remission in IBD, we analyzed publicly available microarray datasets from three independent UC patient cohorts treated with distinct biologic agents. Across multiple therapeutic cohorts, the chemokines CXCL1, CXCL11, and CCL2 were consistently elevated at baseline and demonstrated a striking pattern following treatment (Figure 2B, E, H).

**Figure 1.**
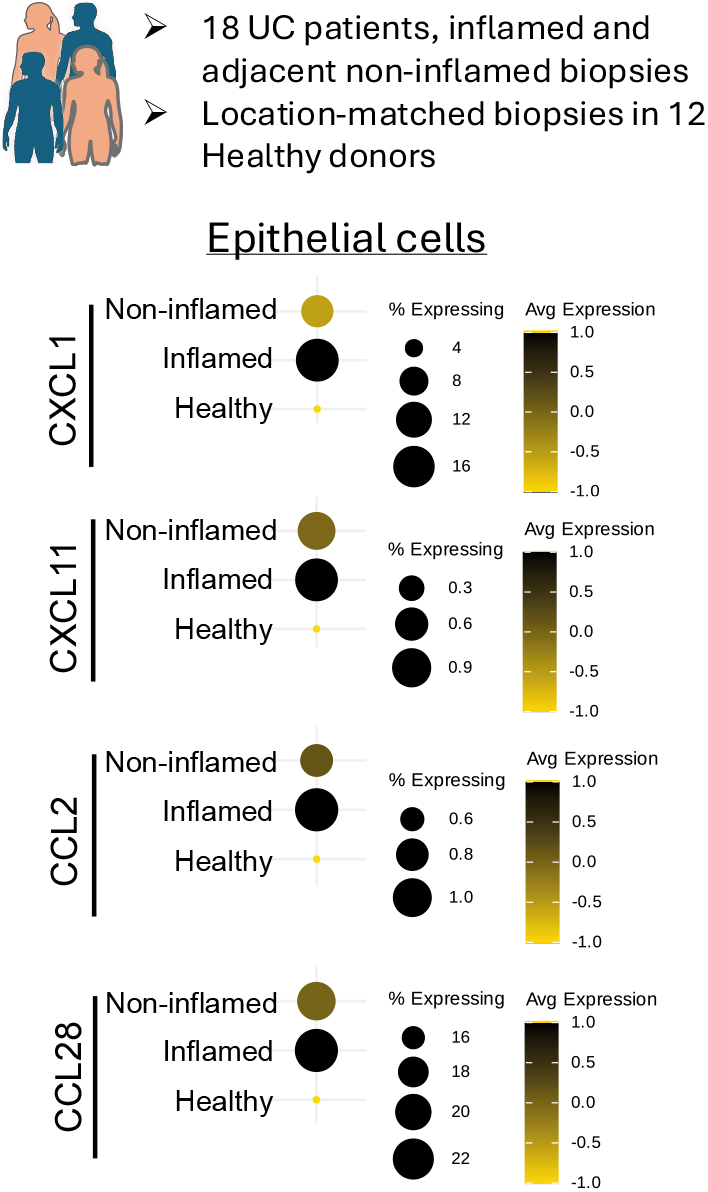
Epithelial-intrinsic chemokine expression in human UC colonoids and tissues. (A) Dot plots from the human colon scRNA-seq data (single cell portal accession=SCP259) showing chemokine expression in epithelial cells from non-inflamed UC, inflamed UC, and healthy donor biopsies.

**Figure 2.**
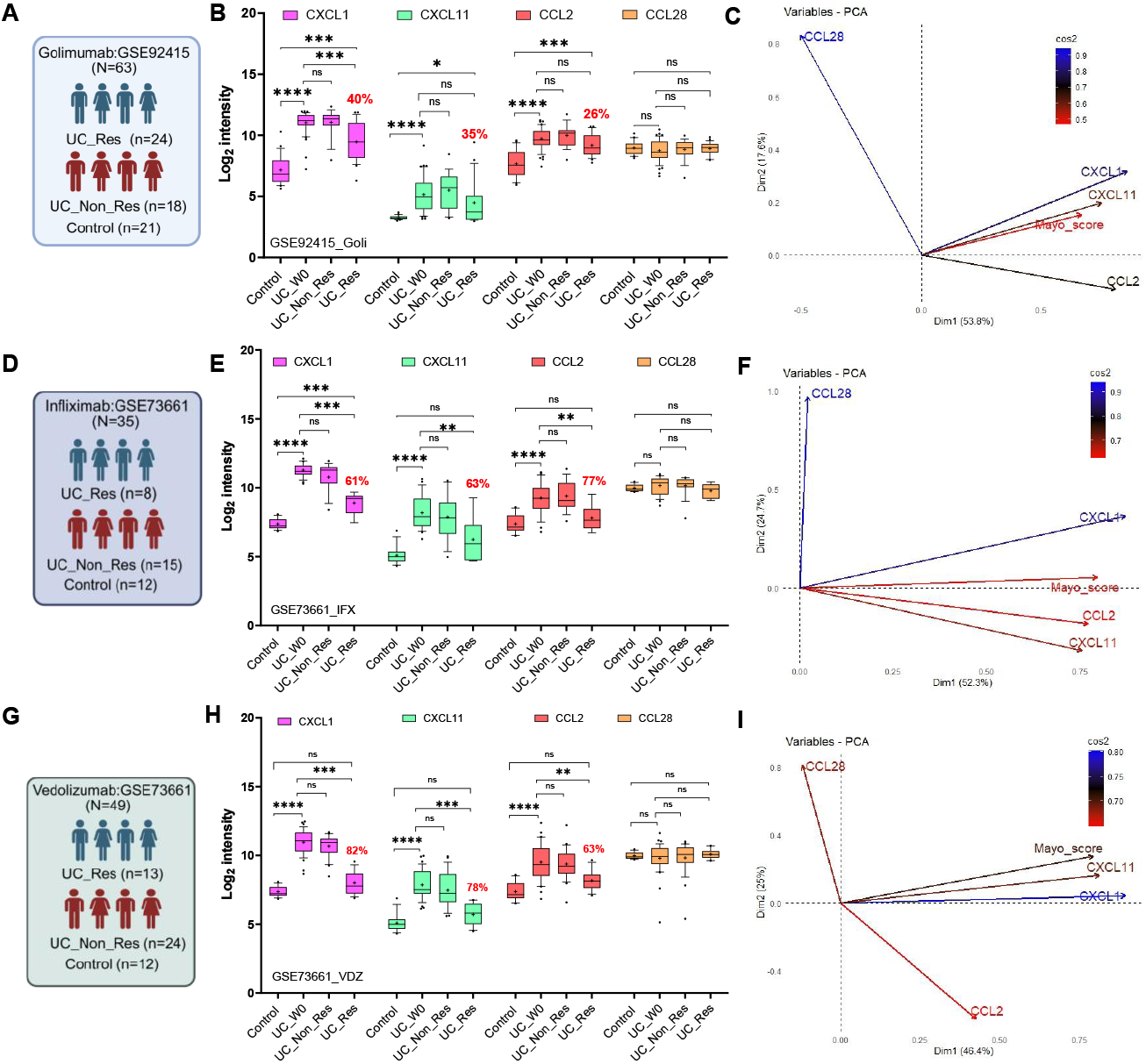
Bulk transcriptomics reveal chemokine expression patterns in UC colonic mucosa across three cohorts before and after biologic treatments. Schematic of the patient cohort are shown in A, D, and G. Box plots (B, E, H) showing Log2 expression intensity of CXCL1, CXCL11, CCL2, and CCL28 in healthy controls (Control), UC patients at baseline prior to treatment (UC_W0), UC non-responders (UC_Non_Res), and UC responders (UC_Res) treated with golimumab (B), infliximab (E), or vedolizumab (H). Percentages indicate the efficacy of the therapy on chemokine expression in UC. *p<0.05, **p<0.001, ***p<0.0001; ns = not significant. (C, F, I) Principal component analysis (PCA) variable plots showing the contribution and correlation of CXCL1, CXCL11, CCL2, CCL28, and Mayo clinical disease activity score to principal components 1 and 2 in the golimumab (C), infliximab (F), and vedolizumab (I) cohorts. Schematics in A, D, and G were prepared in BioRender.

Normalization of chemokine expression occurred only in patients who achieved clinical remission, while non-responders retained elevated expression regardless of therapy type or duration. This pattern was observed across anti-TNF and anti-integrin therapies, suggesting that normalization of epithelial chemokines reflects a shared endpoint of effective treatment rather than a therapy-specific effect. In contrast, CCL28 expression remained relatively stable across disease states and treatment outcomes, indicating that not all epithelial chemokines are dynamically regulated in relation to clinical response (Figure 2B, E, H). Principal component analysis further demonstrated that CXCL1 and CXCL11 strongly tracked with clinical disease activity and contributed to the primary axes of transcriptional variation, whereas CCL28 exhibited a distinct pattern that was not associated with disease severity (Figure 2C, F, I).

To validate our previous findings of upregulated chemokine levels in UC compared to non-IBD control organoids, we analyzed a publicly available scRNA-seq dataset ^10^ containing paired inflamed and non-inflamed colonic biopsies from 18 UC patients and 12 healthy donors.

Epithelial transcriptional profiling of this data revealed a progressive increase in chemokine expression from healthy tissue to non-inflamed UC mucosa and, ultimately, to inflamed mucosa, indicating that epithelial activation precedes overt histologic inflammation and may extend into regions that appear endoscopically quiescent (Figure 1).

To identify the epithelial cellular sources underlying these patterns, we interrogated single-cell RNA sequencing data from the TAURUS-IBD study ^9^, encompassing over 305,862 colonic epithelial cells from patients receiving adalimumab therapy (Figure 3A, 2B). In UC patients, CXCL1, CXCL11, and CCL28 were broadly upregulated across multiple epithelial cell types, especially in organoid-forming progenitor cells such as LGR5-positive stem cells and tuft cells ^11, 12^, suggesting that inflamed epithelial progenitor cells contribute to the chemokine gradient that attracts leukocytes to the damaged epithelium in IBD. Importantly, clinical remission was associated with normalization of CXCL1, CXCL11, and CCL28 expression in stem cells and enterocytes, while non-remission patients maintained elevated expression after therapy (Figure 3C). In contrast, patients with Crohn’s disease (CD) exhibited distinct patterns that are independent of clinical remission (Figure 3D). Following adalimumab treatment, CXCL1, CXCL11, CCL2, and CCL28 showed near-healthy levels in both remission and non-remission groups across most epithelial cell types except in tuft cells (Figure 3D), suggesting that epithelial chemokine programs are uncoupled from clinical response in Crohn’s disease.

**Figure 3.**
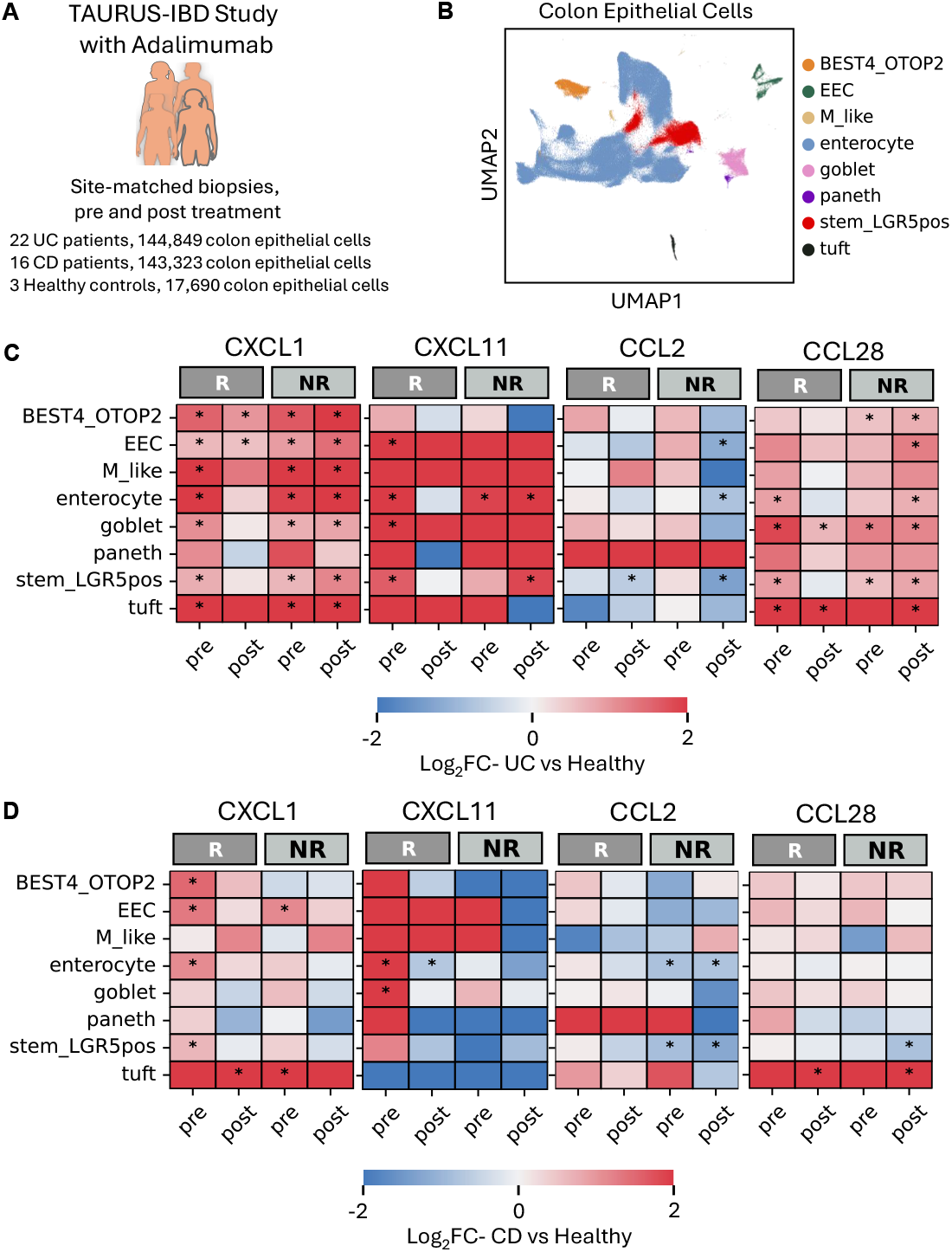
Stem cell-derived chemokines track clinical remission in IBD in a disease-specific manner. (A) Schematic of the TAURUS-IBD study design. (B) Uniform Manifold Approximation and Projection (UMAP) of 305,862 colonic epithelial cells colored by cell type. (C-D) Heatmaps showing log2 fold change (Log2 FC) of CXCL1, CXCL11, CCL2, and CCL28 expression relative to healthy controls across epithelial cell subtypes in UC (C) and CD (D) patients. Columns represent remission (R) and non-remission (NR) patients before (pre) and after (post) adalimumab treatment. Asterisks denote FDR-adjusted p < 0.05 and |Log_2_ FC| ≥ 0.5 compared to healthy control cells.

## 3. Discussion

Chemokine-directed leukocyte trafficking to the sites of injury is a hallmark of inflammation in IBD ^3^. Recently, the overexpression of CXCL1, CXCL11, CCL2, and CCL28 was reported in colon epithelial organoids derived from UC patients ^6^. In this report, we used bulk and single-cell transcriptomic data from 4 clinical trials with anti-TNFα and anti-integrin therapies to investigate whether chemokines retained in UC epithelial organoids improved in patients who achieved clinical remission after treatment. Bulk transcriptomic analyses showed a consistent inverse relationship between epithelial chemokines, especially CXCL1 and CXCL11, and clinical remission in UC patients for all evaluated therapies. In the UC colon, we observed cell-type-specific resolution of epithelial CXCL1, CXCL11, and CCL28, particularly in progenitor cells, where epithelial chemokine normalization distinguished responders from non-responders in UC, but not in Crohn’s. These data demonstrate that chemokines retained in UC organoids serve as biomarkers of treatment response in UC but not in CD, highlighting disease-specific differences that can be functionally targeted in organoid models.

The upregulation of these chemokines in organoid-forming progenitor cells, such as LGR5-positive stem cells and tuft cells ^11, 12^ suggests that inflamed epithelial progenitor cells contribute to the chemokine gradient that attracts leukocytes to the damaged epithelium in IBD. The previously reported chemokine overexpression in active UC organoids is only prominent during differentiation ^6^, and agrees with the concept of inflammation memory in the stem-cell compartment ^13^, indicating that the activity of inflamed progenitor-like cells may be tied to chemokine overexpression in UC. Importantly, our current data analyses showed that this epithelial progenitor-encoded inflammatory memory is only erased upon effective therapeutic intervention, highlighting that successful therapy must ultimately reset epithelial programs at their source rather than solely suppress downstream immune responses. Since only 27% of UC patients in the TAURUS-IBD study achieved remission ^9^, candidate therapies targeting chemokine disparities retained in patient-derived IBD epithelial organoids may increase the likelihood of chemokine normalization in stem cells, reduce leukocyte trafficking to the epithelium, and improve the odds of clinical remission in UC.

The divergence observed between UC and CD provides additional insight into disease-specific mechanisms, especially for epithelial-immune interactions. In UC, epithelial chemokine expression appears tightly coupled to clinical outcomes, implying a direct relationship between epithelial state and disease activity. In contrast, the normalization of chemokine expression in CD, regardless of clinical response, suggests that epithelial pathways for the analyzed chemokines may be less central to disease persistence, or that other factors, such as immune compartment dynamics, may play a more dominant role. This distinction may have important implications for therapeutic development, as it suggests that epithelial-targeted strategies may be particularly relevant in UC. As our results are from colon-derived data, it is also plausible that samples from other intestinal locations in CD may exhibit similar or distinct epithelial chemokine signatures.

Overall, our analyses demonstrate that epithelial chemokines in organoid-forming progenitor cells, particularly CXCL1, CXCL11, and CCL28, track clinical disease activity and resolve only in patients who achieve remission following biologic therapy. These chemokines distinguish responders from non-responders in UC but not in CD, underscoring disease-specific epithelial-immune interactions in the colon. The retention of this inflammatory chemokine signature in patient-derived UC organoids validates these chemokines as functional readouts for epithelial-directed therapeutic screening using patient-derived organoid models.

## 4. Acknowledgments

This work was supported by the National Institute of Diabetes and Digestive and Kidney Diseases of the National Institutes of Health (NIH) under Award Numbers R00DK136971 to B.A.O. The content is solely the responsibility of the authors and does not necessarily represent the official views of the NIH. The study was also supported by the UAB IMPACT funds, UAB Workforce Empowerment Award, and the UAB Kaul Pediatric Research Award to B.A.O. This work utilized Cheaha computing resources provided by the UAB IT’s Research Computing Group.

